# Short-term aerobic training does not improve memory functioning in relapsing remitting multiple sclerosis – a randomized controlled trial

**DOI:** 10.1101/366161

**Authors:** L Baquet, H Hasselmann, S Patra, JP Stellmann, E Vettorazzi, AK Engel, S Rosenkranz, J Poettgen, SM Gold, KH Schulz, C Heesen

## Abstract

**Background:** Only few aerobic exercise intervention trials specifically targeting cognitive functioning have been performed in MS.

**Objective and methods:** This randomized controlled trial aimed to determine the effects of aerobic exercise on cognition in relapsing-remitting MS. The primary outcome was verbal memory (Verbal learning and memory test, VLMT). Patients were randomized to an intervention group (IG) program or a waitlist control group (CG). Patients in the IG exercised according to an individually tailored training schedule (with 2-3 sessions per week for 12 weeks). The primary analysis was carried out using the intention-to-treat (ITT) sample with ANCOVA adjusting for baseline scores.

**Results:** 77 RRMS patients were screened and 68 participants randomized (CG n=34; IG n=34). The sample comprised 68% females, had a mean age of 39 years, a mean disease duration of 6.3 years, and a mean EDSS of 1.8. No significant effects were detected in the ITT analysis for the primary endpoint VLMT or any other cognitive measures. Moreover, no significant treatment effects were observed for quality of life, fatigue, or depressive symptoms.

**Conclusion:** This study failed to demonstrate beneficial effects of aerobic exercise on cognition in RRMS.

The trial was prospectively registered at clinicaltrials.gov (NCT02005237).

## Introduction

While the pathological hallmark of multiple sclerosis (MS) is inflammation and demyelination of the central nervous system, progressive neurodegeneration plays an important role in MS pathobiology and occurs early on in the disease process (Reich et al. 2018). So-called disease modifying therapies can control the inflammation to a variable extent and are thus able to decrease the frequency of relapses (Wingerchuk & Weinshenker 2016). However, there is only modest evidence that the long-term disease progression is altered. Neurodegeneration is thought to be a major driver of MS-related impairments in higher order brain functions such as cognition, highlighting the importance of developing treatment options in this area (Amato et al. 2013).

Exercise training has received increasing attention as a putative strategy to target the degenerative component of multiple sclerosis (Motl et al. 2017). Some (Kim & Sung 2017), albeit not all (Klaren et al. 2016) studies conducted in experimental autoimmune encephalomyelitis (the animal model of MS) have indicated a neuroregenerative effect of exercise. For example, Kim et al. showed that exercise can improve memory function and increase hippocampal neurogenesis in experimental autoimmune encephalomyelitis (EAE) (Kim & Sung 2017).

Moreover, there are clinical data that support the neuroprotective potential of exercise. In healthy elderly humans, aerobic exercise training selectively increased the size of the anterior hippocampus and change in hippocampal volume was significantly related to improved memory in the exercise group (Erickson et al. 2011). Epidemiological studies have reported an association of physical fitness with lower risk of dementia disorders (Winblad et al. 2016). In addition, lifestyle interventions including exercise and nutrition counselling can reduce the risk of cognitive deterioration in people with minimal cognitive deficits (Ngandu et al.). Thus, cognitive performance seems to be a suitable endpoint to determine neuroprotective effects of exercise interventions.

However, while numerous studies now have shown beneficial effects of exercise training on physical capacity, strength, quality of life, mood (Latimer-Cheung et al. 2013) and fatigue in MS (Heine et al. 2015), high quality clinical trials focusing on cognitive function are scarce and the evidence not conclusive (Sandroff et al. 2016). To our knowledge no study yet has addressed cognition as its primary endpoint.

In the current study, we aimed to determine the effects of individually tailored aerobic exercise on verbal learning and memory as the primary endpoint in patients with relapsing-remitting MS.

## Material & Methods

### Study design, overview and patient recruitment

Our trial was a single-blind, 1:1 randomized, controlled phase IIa study with an active intervention (IG) and a waitlist control group (CG). Patients in the IG underwent bicycle ergometry training with 2-3 sessions per week for 12 weeks and a subsequent extension phase of another 12 weeks. During the extension phase, patients originally randomized to the waitlist group had the opportunity to train as well and intervention group patients were invited to continue training. Details regarding the intervention are provided below.

The primary endpoint of our study was the Verbal Learning and Memory Test (VLMT) (Lux et al. 1999). Secondary endpoints were other cognitive functions (see below for details), as well as neuroimaging parameters (which will be reported separately). Tertiary endpoints were walking ability and motor function (for details see below) as well as patient reported outcomes (for details see below).

All endpoints were obtained at the MS Day Hospital at the University Medical Center Hamburg Eppendorf at baseline (t0), after the intervention (week 12; t1) and at the end of the extension phase at week 24 (t2). Patient recruitment was conducted by screening the registry database of the MS Day Hospital at the University Medical Center Hamburg Eppendorf for patients who met inclusion criteria and had indicated an interest in information about new clinical studies. Recruitment started in January 2013 and was completed in November 2015.

### Standard protocol approvals and patient consent

The trial was approved by the ethics committee of the Hamburg Chamber of Physicians, (Registration Number PV4356). Participants gave written consent before enrolling in the study. The study was prospectively registered at ClinicalTrials.gov (identifier NCT02005237).

### Inclusion and exclusion criteria

To be eligible for trial participation, patients had to be diagnosed with relapsing remitting MS (RR-MS) according to the McDonald criteria 2010 (Polman et al. 2011), an EDSS score <3.5, and currently in remission with no relapse or progression during the last 3 months. Patients had to be on stable immunotherapy for more than three months or without any planned treatment for the next 6 months.

Exclusion criteria were severe cognitive impairment or major psychiatric comorbidity (e.g. severe depression, psychosis, dementia) based on clinical judgement. Patients not capable to undergo aerobic exercise for medical reasons (i.e, due to heart disease) were excluded.

### Sample size and randomization

Sample size calculation was based on the results obtained in our previous trial (Briken et al. 2014). For the VLMT learning subtest we calculated group sample sizes of 20 each to achieve 90% power to detect a difference of −5.8 points on the scale. For the VLMT delayed recall we calculated group sample sizes of 17 and 17 to achieve 92% power to detect a difference of −2.6. Significance level (alpha) was set as 0.025 using a two-sided two-sample t-test. For the dual primary endpoints 90% of power could thus be achieved by recruiting n=20 patients for each arm of the trial. To safeguard against dropouts and the possibility that the effect sizes in the pilot trial had overestimated the real effect, we planned to enroll at least n=60 patients (approximately n=30 in each trial arm). After reaching the intended sample size the trial was stopped.

Inclusion was performed by physicians from the MS Day Hospital. Random sequence was generated by E. Vettorazzi. Using a computer-based random number generator, we performed randomization by concealed allocation. While blinding of the participants was not possible, the outcome measures were obtained by assessors blind to group assignment.

### Intervention

The intervention comprised 12 weeks of aerobic exercise on a bicycle ergometer. The pre-specified interval training schedule was tailored to the patients’ individual level of aerobic fitness as determined by a spiroergometric exhaustion test at baseline (see trial protocol for details). The patients had to train 2-3 times a week for 12 weeks. Duration and power (in watts) in each training session were gradually increased to achieve a continuous increase of performance while keeping the perceived (Borg scale) and objectively measured (heart rate) exhaustion at a constant level. Each session consisted of 3-5 intervals with short breaks and lasted for a maximum of 70 minutes each. Details are provided in the trial protocol (see Suppl. material). Experienced physiotherapists who were familiar with the training program supervised the training.

### Outcome measures

#### Cognitive functions

From the Verbal Learning and Memory Test (VLMT) (Lux et al. 1999) we used the learning curve (words memorized in trials 1-5) and the delayed recall as a measure of memory. This choice was based on several considerations: 1. While processing speed is often considered the most sensitive screening test for cognitive impairment in MS (Costa et al. 2016), impairments in learning and memory are also highly common in this population (Chiaravalloti & DeLuca 2008); 2. Basic science and clinical trials in healthy aging have indicated that CNS networks involved in learning and memory are particularly affected by exercise (Loprinzi et al. 2017) and 3. Results from our previous trial in progressive MS suggested that verbal learning and memory was most strongly improved by exercise.

Other neuropsychological functions were tested as secondary endpoints. We chose the Symbol-Digit-Modalities Test (SDMT) and the Paced Auditory Serial Addition Test (PASAT) to assess information processing (Fischer et al. 1999) (López-Góngora et al. 2015). Visuospatial learning and memory was assessed using the Brief Visuospatial Memory Test - Revised (BVMT-R) (Tam & Schmitter-Edgecombe 2013). In addition, we applied the Corsi Block-Tapping Task (Berch et al. 1998) to assess short term and working memory. Three subtests of the Test Battery for Attention (TAP) were used to assess attention – alertness, covert shift of attention (CSA) and incompatibility (Zimmermann & Fimm 1994). We administered the Regensburg Verbal Fluency Test (RWT) as a measure of verbal fluency (Aschenbrenner et al. 2000). To assess social cognition / theory of mind, we utilized an abbreviated version of the Movie for the Assessment of Social Cognition (MASC) (Pöttgen et al. 2013).

#### Motor function & Walking Ability

The Six Minute Walking Test (6MWT) (Goldman et al. 2008) and the Timed 25-Foot Walk (T25FW) (Kaufman et al. 2000) served to assess walking ability. The Nine-Hole Peg Test (9HPT) assessed motor function of the upper body (Feys et al. 2017).

#### Patient-reported outcome measures

We included the 16-item Inventory of Depressive Symptomatology (IDS16SR) (Fischer et al. 2015), the Fatigue Scale for Motor and Cognitive Function (FSMC) (Penner et al. 2009), the Hamburg Quality of Life Questionnaire for Multiple Sclerosis (HAQUAMS) (Gold et al. 2001) and the Multiple Sclerosis Walking Scale-12 (MSWS-12) (Hobart et al. 2003).

#### Aerobic fitness

Bicycle spiroergometry using a Metalyser 3b (CORTEX Biophysik GmbH, Leipzig, Germany) was performed at baseline, after the intervention (week 12), and at the end of the extension phase (week 24). Peak oxygen intake (VO_2peak_), oxygen intake per kilogram of body weight (VO_2peak_/kg) and maximal power (P_max_) as well as Borg scaling and lactate levels were obtained every two minutes. The test started with 10 watts after a two-minute resting and a two-minute warm-up phase at 10 watts. Performance was continuously increased by 1 watt every 6 seconds which resulted in a ramp of 20 watts/2 min. We obtained continuous recordings of stress ECG to monitor cardiovascular function. Patients were instructed to exercise until perceived exhaustion (Borg 18-20). Spiroergometry results at baseline provided the anchor for the individualized training schedule which was determined by an algorithm including several maximal and sub-maximal parameters of the spiroergometry session (Power, oxygen intake (VO_2_), Rating of Perceived Exertion Scale (Borg scaling), lactate thresholds, ventilatory threshold, Karvonen-Formula, Respiratory Exchange Rate (RER)). Training intervals were performed near the aerobic threshold level within an individually determined starting point to improve aerobic capacity (Westhoff et al. 2013). For training plan details see protocol in the suppl. file.

### Statistical analyses

The primary analysis compared changes from baseline to week 12 between the intervention group and the waitlist control group. According to guidelines for statistical analysis of clinical trials published by The European Agency for the Evaluation of Medicinal Products (CPMP/ICH/363/96 and CPMP/EWP/2863/99), we computed the primary statistical analysis for all outcomes using ANCOVA models adjusting for baseline measurements of the respective outcome variable to evaluate treatment effects (measured as change from baseline). No other covariates were included in this primary analysis. As recommended, this model did also not include treatment by covariate interactions. All primary analyses were conducted as intention-to-treat (ITT) including all patients who had received group allocation. Every effort was made to obtain week 12 and week 24 data from all participants (even if they dropped out of the exercise program). In case of missing data, primary ITT analyses were conducted using a Last-Observation-Carried-Forward (LOCF) approach. As sensitivity analyses, we computed the same ANCOVA models as per-protocol (PP, i.e. including only patients who completed the 12 weeks of training with at least 18 training sessions). We also defined a responder group (RG), which included those patients in the IG who showed the strongest increases (upper tercile, n=11) in P_max_ values from baseline to week 12 (P_max__week12 – P_max__baseline). This group was compared to all patients in the control group in additional sensitivity analyses.

## Results

77 patients were interested in participating in our study. After screening for eligibility, 68 patients fulfilled the inclusion criteria (for patient attrition see Fig. 1). They were randomized to the IG or a CG. On average, patients in the IG received 19.5 training sessions. Notably, the proportion of men was higher in the IG than in the CG (38% versus 26%). 58 patients had complete data at baseline-as well as the follow-up-testing 12 weeks later. For the ITT analyses, all patients randomized were included (i.e. 34 patients in each group).

**Figure 1:**
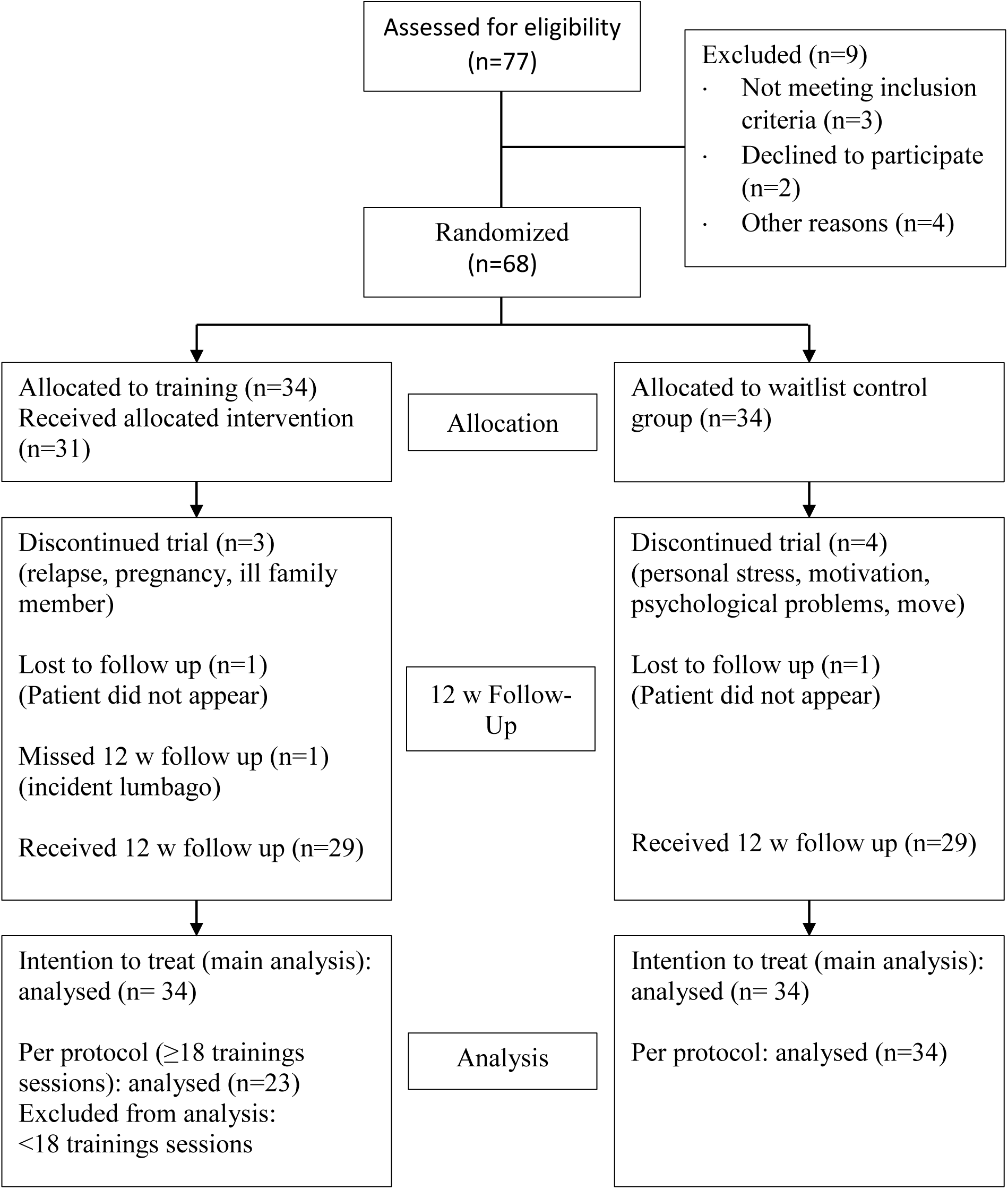
Participant flow chart

Overall, patients had low to moderate clinical disability (median EDSS=1.5; range 0 – 3.5). Cognitive function overall was in the range of what would be expected in early RRMS and as a group only showed mild impairment (PASAT showed the highest rate of individual abnormal values with baseline findings below age and education adapted references in n=5 IG and n=4 CG patients).

### Effects of exercise on cognitive function

No significant treatment effects were detected for the primary endpoints (VLMT learning, VLMT delayed recall) or any of the other neuropsychological tests included. This was seen in the ITT analyses as well as the sensitivity analyses (PP and responder analyses).

### Effects of exercise on motor function and walking ability

Similarly, ITT analyses of motor function was negative with no significant effects in 3 measures (6MWT, T25, 9HPT dominant hand). One test (9HPT non-dominant hand) showed significant effects in favor of the control group (see table 3).

**Table 1:** Clinical baseline characteristics

|  | CG<br>Baseline<br>n=34 |  | IG_ITT<br>Baseline<br>n=34 |  | IG_PP<br>Baseline<br>n=23 |  | IG_RG<br>Baseline<br>n=11 |  |
| --- | --- | --- | --- | --- | --- | --- | --- | --- |
| Age (years) | 39.6 | (9.7) | 38.2 | (9.6) | 38.6 | (9.9) | 38.2 | (9.6) |
| Sex (m/f) | 9/25 |  | 13/21 |  | 9/14 |  | 5/6 |  |
| Education (years) | 12.1 | (1.5) | 12.3 | (1.4) | 12.7 | (1.0) | 12.1 | (1.6) |
| EDSS Baseline | 1.8 | (1.0) | 1.7 | (0.9) | 1.6 | (0.9) | 1.4 | (0.9) |
| Disease duration - symptoms (years) | 9.1 | (7.7) | 8.1 | (5.7) | 7.2 | (6.2) | 5.8 | (6.7) |
| Disease duration - diagnosis (years) | 5.7 | (6.3) | 6.8 | (5.5) | 6.6 | (6.0) | 4.4 | (6.6) |
| Immunotherapy (%) | 44.1 |  | 67.7 |  | 69.6 |  | 72.7 |  |
| Number of sessions |  |  | 19.5 | (10.0) | 24.7 | (6.5) | 26.1 | (7.0) |

Data as mean (standard deviation)
CG: Control group; IG: Intervention group; ITT: intention to treat; PP: per protocol; RG: Responder subgroup EDSS: Expanded Disability Status Scale

**Table 2:**
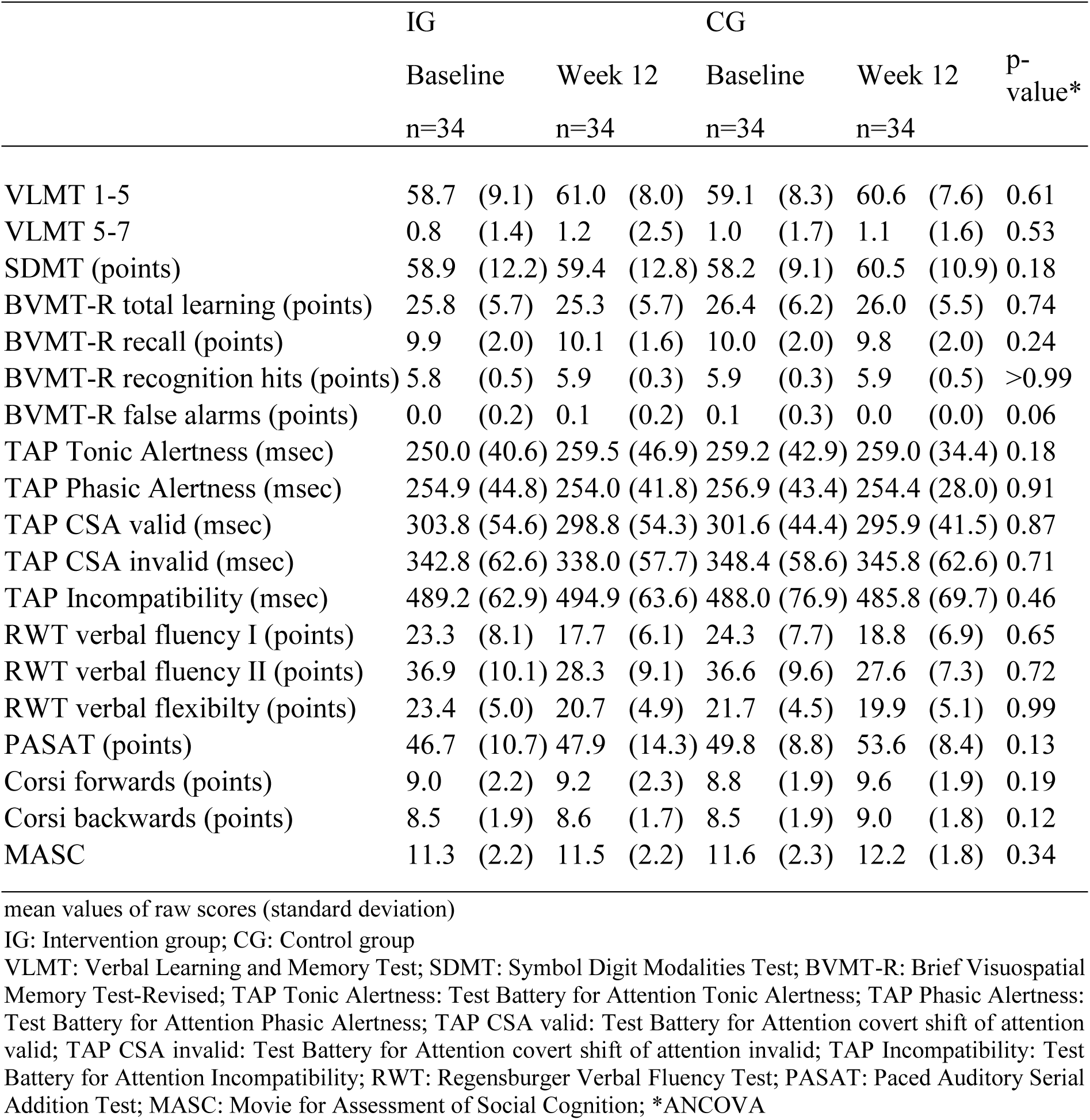
Cognitive outcomes

|  | IG |  |  |  | CG |  |  |  | p-value* |
| --- | --- | --- | --- | --- | --- | --- | --- | --- | --- |
|  | Baseline |  | Week 12 |  | Baseline |  | Week 12 |  |  |
|  | n=34 |  | n=34 |  | n=34 |  | n=34 |  |  |
| VLMT 1-5 | 58.7 | (9.1) | 61.0 | (8.0) | 59.1 | (8.3) | 60.6 | (7.6) | 0.61 |
| VLMT 5-7 | 0.8 | (1.4) | 1.2 | (2.5) | 1.0 | (1.7) | 1.1 | (1.6) | 0.53 |
| SDMT (points) | 58.9 | (12.2) | 59.4 | (12.8) | 58.2 | (9.1) | 60.5 | (10.9) | 0.18 |
| BVMT-R total learning (points) | 25.8 | (5.7) | 25.3 | (5.7) | 26.4 | (6.2) | 26.0 | (5.5) | 0.74 |
| BVMT-R recall (points) | 9.9 | (2.0) | 10.1 | (1.6) | 10.0 | (2.0) | 9.8 | (2.0) | 0.24 |
| BVMT-R recognition hits (points) | 5.8 | (0.5) | 5.9 | (0.3) | 5.9 | (0.3) | 5.9 | (0.5) | >0.99 |
| BVMT-R false alarms (points) | 0.0 | (0.2) | 0.1 | (0.2) | 0.1 | (0.3) | 0.0 | (0.0) | 0.06 |
| TAP Tonic Alertness (msec) | 250.0 | (40.6) | 259.5 | (46.9) | 259.2 | (42.9) | 259.0 | (34.4) | 0.18 |
| TAP Phasic Alertness (msec) | 254.9 | (44.8) | 254.0 | (41.8) | 256.9 | (43.4) | 254.4 | (28.0) | 0.91 |
| TAP CSA valid (msec) | 303.8 | (54.6) | 298.8 | (54.3) | 301.6 | (44.4) | 295.9 | (41.5) | 0.87 |
| TAP CSA invalid (msec) | 342.8 | (62.6) | 338.0 | (57.7) | 348.4 | (58.6) | 345.8 | (62.6) | 0.71 |
| TAP Incompatibility (msec) | 489.2 | (62.9) | 494.9 | (63.6) | 488.0 | (76.9) | 485.8 | (69.7) | 0.46 |
| RWT verbal fluency I (points) | 23.3 | (8.1) | 17.7 | (6.1) | 24.3 | (7.7) | 18.8 | (6.9) | 0.65 |
| RWT verbal fluency II (points) | 36.9 | (10.1) | 28.3 | (9.1) | 36.6 | (9.6) | 27.6 | (7.3) | 0.72 |
| RWT verbal flexibility (points) | 23.4 | (5.0) | 20.7 | (4.9) | 21.7 | (4.5) | 19.9 | (5.1) | 0.99 |
| PASAT (points) | 46.7 | (10.7) | 47.9 | (14.3) | 49.8 | (8.8) | 53.6 | (8.4) | 0.13 |
| Corsi forwards (points) | 9.0 | (2.2) | 9.2 | (2.3) | 8.8 | (1.9) | 9.6 | (1.9) | 0.19 |
| Corsi backwards (points) | 8.5 | (1.9) | 8.6 | (1.7) | 8.5 | (1.9) | 9.0 | (1.8) | 0.12 |
| MASC | 11.3 | (2.2) | 11.5 | (2.2) | 11.6 | (2.3) | 12.2 | (1.8) | 0.34 |

mean values of raw scores (standard deviation)
IG: Intervention group; CG: Control group
VLMT: Verbal Learning and Memory Test; SDMT: Symbol Digit Modalities Test; BVMT-R: Brief Visuospatial Memory Test-Revised; TAP Tonic Alertness: Test Battery for Attention Tonic Alertness; TAP Phasic Alertness: Test Battery for Attention Phasic Alertness; TAP CSA valid: Test Battery for Attention covert shift of attention valid; TAP CSA invalid: Test Battery for Attention covert shift of attention invalid; TAP Incompatibility: Test Battery for Attention Incompatibility; RWT: Regensburger Verbal Fluency Test; PASAT: Paced Auditory Serial Addition Test; MASC: Movie for Assessment of Social Cognition; \*ANCOVA

**Table 3:** Motor, training and patient reported outcomes

|  | IG |  |  |  | CG |  |  |  | P-value* |
| --- | --- | --- | --- | --- | --- | --- | --- | --- | --- |
|  | Baseline |  | Week 12 |  | Baseline |  | Week 12 |  |  |
|  | n=34 |  | n=34 |  | n=34 |  | n=34 |  |  |
| Motor function and aerobic fitness |  |  |  |  |  |  |  |  |  |
| 6 MWT (m) | 432.1 | (105.7) | 454.4 | (104.1) | 448.8 | (79.8) | 466.7 | (84.2) | 0.85 |
| 9 HPT dominant (sec) | 19.5 | (3.7) | 19.6 | (3.6) | 19.1 | (2.9) | 18.6 | (3.0) | 0.11 |
| 9 HPT non dominant (sec) | 19.6 | (2.9) | 19.9 | (3.0) | 19.8 | (4.1) | 19.0 | (3.5) | 0.03 |
| T25FW (sec) | 4.8 | (0.8) | 4.8 | (0.8) | 4.8 | (0.8) | 4.8 | (0.8) | 0.49 |
| V<sub>O<sub>2</sub>peak</sub> (ml O<sub>2</sub>/min) | 2151.5 | (669.5) | 2225.6 | (743.4) | 1761.5 | (421.3) | 1779.4 | (427.4) | 0.37 |
| V<sub>O<sub>2</sub>peak</sub>/kg ((ml O<sub>2</sub>/min)/kg) | 27.2 | (7.9) | 28.0 | (8.8) | 25.6 | (5.5) | 25.6 | (5.4) | 0.24 |
| P<sub>max</sub> (watt) | 160.4 | (45.4) | 176.3 | (55.4) | 139.5 | (31.1) | 139.6 | (31.0) | 0.01 |
| Patient-reported outcome measures |  |  |  |  |  |  |  |  |  |
| IDS-16SR | 6.4 | (5.1) | 5.9 | (4.4) | 6.1 | (4.3) | 6.3 | (4.6) | 0.44 |
| FSMC | 51.5 | (22.1) | 51.1 | (22.6) | 53.4 | (21.6) | 50.9 | (21.4) | 0.44 |
| MSWS-12 | 19.4 | (10.4) | 19.6 | (10.2) | 18.7 | (10.7) | 18.7 | (9.9) | 0.78 |
| HAQUAMS | 53.1 | (16.9) | 53.7 | (17.8) | 51.2 | (18.7) | 51.8 | (14.7) | 0.75 |

Data as mean (standard deviation)
IG: Intervention group; CG: Control group
6 MWT: Six minute walking test; 9 HPT: Nine-Hole Peg Test; T25FW: Timed 25-foot walk; P<sub>max</sub>: maximal Power;
IDS16-SR: 16-item version of Inventory of Depressive Symptomatology Self-Rated; FSMC: Fatigue Scale for Motor and Cognitive Functions; MSWS-12: 12-item MS Walking Scale; HAQUAMS: Hamburger Quality of Life Questionnaire in Multiple Sclerosis; \*ANCOVA

### Effects of exercise on fitness

Significant treatment effects favoring the intervention group were observed for P_max_ in the ITT analysis (see Table 3). This was confirmed in the PP analysis. However, VO_2_ Peak and VO_2_/kg showed no significant treatment effects.

Reflecting on baseline fitness indices with 140-160 watts at P_max_ and 25-27 ml (O_2_/min)/kg at V0_2peak_, the sample showed a deficient fitness level.

### Effects of exercise on patient-reported outcomes

Finally, there were no significant treatment effects in any of the patient-reported outcome measures including quality of life, fatigue, mood or self-reported walking ability (see table 3).

## Discussion

Overall, our trial of standardized exercise over 12 weeks in RRMS patients compared to a waitlist control group failed to meet its clinical endpoints. Specifically, the study did not show effects on any cognitive measure and failed to produce significant changes in any of the obtained patient related or objective outcome measures for functioning in MS. These results thus cast doubt on the generalizability of recent trials that suggested the potential of exercise to improve cognitive function in progressive (Briken et al. 2014) or relapsing-remitting MS (Zimmer et al. 2017).

Compared to our previous study in progressive MS patients (Briken et al. 2014), RRMS patients in the current trial were substantially less disabled physically and cognitively, thereby potentially limiting room for improvements (ceiling effect). Mean EDSS values were below 2.0 which were the lowest among 26 recently reviewed exercise interventions in MS assessing cognition as an outcome (Sandroff et al. 2016). In addition, all mean baseline scores on cognitive function were within 1 SD of normative values. In contrast, unselected RRMS cohorts show cognitive deficits in at least 40% (Chiaravalloti & DeLuca 2008). Interestingly from the 26 studies referred to above, only two case report studies selected patients based on cognitive deficits. Our study underlines that further studies on the cognitive effects of exercise treatments need to address patients presenting with relevant cognitive deficit. Another aspect to consider is the choice of verbal learning and memory as our primary endpoint. As described in the Methods section above, this choice was based on several considerations. However, it could be argued that – given its high sensitivity for detecting cognitive impairment in MS – a test of processing speed (e.g. the SDMT or the PASAT) might have been a more suitable choice. However, given our null effects across all cognitive domains tested as primary or secondary endpoints (including the SDMT and PASAT), this would not have changed our results.

One strength of our study is that cognition was predefined as the primary outcome. Moreover, based on our previous work we chose verbal learning and memory as key domain. But is verbal learning and memory the neurocognitive domain most sensitive to exercise interventions? Consistent with our earlier study in progressive MS, a recent trial on high intensity training could show effects on different cognitive dimensions with strongest effects in the verbal memory dimension of the BICAMS (Zimmer et al. 2017). However, in the few other MS exercise studies most effects were reported in information processing speed and executive functioning (Sandroff et al. 2016). From studies on healthy aging it appeared that depending on age and intervention different cognitive domains might be influenced (Hötting & Röder 2013). For example, Hötting et al. have shown an improvement of episodic memory in middle-aged adults after cardiovascular training but enhanced attention scores after a combined stretching/coordination training (Hötting et al. 2012). Thus, further work is needed to address which domains might be most sensitive for which intervention at which intensity and at which disease stage in MS.

Training intensity – albeit carefully tailored to the individual patient’s level of fitness at baseline - was only moderate. While available meta-analyses indicate a consistent benefit of exercise training in MS on fitness (Latimer-Cheung et al. 2013), this effect can be subtle or not detectable in individual trials. In our previous trial in progressive MS (Briken et al. 2014) with a similar number of training sessions over 10 weeks, VO_2_ peak increases in the IG were small and between group comparisons reached significance mainly due to worsening of the control group. An earlier study in a mixed sample of relapsing and progressive patients of 8 weeks duration failed to detect significant increases in VO_2peak_ (Schulz et al. 2004). Therefore, a more challenging training plan might be considered in future trials, especially in patients with minimal neurological impairment.

Having said that, effects of exercise training on cognitive functioning in other populations seem not to directly depend on fitness parameters. Resistance training or coordinative training regimen have been shown to improve different cognitive measures (Hötting & Röder 2013). Therefore, cardiovascular fitness indicators can only be regarded as surrogate markers and not mandatory outcomes for beneficial effects on brain functioning.

In a related matter, training duration might have been too short. MS exercise studies to date typically employ training programs of approximately 12 week duration (Latimer-Cheung et al. 2013) with the longest studies up to 26 weeks. Studies in ageing healthy adults performed training in up to 52 weeks and meta-analytic data indicate a relevant effect of intervention length (Northey et al. 2017). Intriguingly, the recent study by Zimmer indicated that high intensity training over just 3 weeks can have beneficial effects on cognition as well other functional domains (Zimmer et al. 2017). Therefore, an optimal balance of intensity and duration needs to be achieved and weighted against other factors such as treatment adherence and attrition. In this context, acceptance of a control condition needs to be discussed.

Finally, on a more optimistic note, it is conceivable that central nervous system (CNS) effects of exercise occur on a subclinical level, i.e. without detectable effects on formal cognitive testing. For example, we recently showed that resistance training can alter structural MRI markers in MS (Kjølhede et al. 2017). Thus, MRI might be a more suitable tool to detect early alterations of functional and structural brain status, particularly in short trials. In the current trial, we have obtained both measures of structural and functional connectivity, which will be reported separately.

## Conclusion

In conclusion we could not demonstrate a beneficial effect of a 12 week moderate exercise training in minor disabled MS on a set of cognitive outcome measures. This negative finding can help to design further exercise trials in MS.

